# The mammalian SKI complex is a broad-spectrum antiviral drug target that upregulates cellular cholesterol to inhibit viral replication

**DOI:** 10.1101/2024.12.03.626536

**Authors:** Stuart Weston, Lauren Baracco, Louis Taylor, Alison Scott, Gaurav Kumar, Paul Shapiro, Alexander D. MacKerell, Matthew B. Frieman

## Abstract

There is a need for the development of broad-spectrum antiviral compounds that can act as first line therapeutic countermeasures to emerging viral infections. Host-directed approaches present a promising avenue of development and carry the benefit of mitigating risks of viral escape mutants. We have previously found the SKI (super killer) complex to be a broad-spectrum, host-target with our lead compound ("UMB18") showing activity against influenza A virus, coronaviruses, and filoviruses. The SKI complex is a cytosolic RNA helicase and we previously found that UMB18 inhibited viral RNA production but did not further define the mechanism. Here, we demonstrate that UMB18 directly binds to SKIC8 of the SKI complex and through transcriptomic analysis of UMB18 treated A549 cells revealed an upregulation of genes in the mevalonate pathway which drives cholesterol synthesis. Further investigation validated the genetic upregulation and confirmed an increase in total cellular cholesterol. This upregulation was dependent on the sterol regulatory element binding proteins (SREBPs) and their regulator SCAP, the major regulators for cholesterol and fatty acid synthesis. Depletion of the SREBPs or SCAP with siRNA, or extraction of cholesterol with methyl β-cyclodextrin attenuated UMB18 antiviral activity, emphasizing the role of increased cholesterol synthesis in this mechanism of action. Our findings further define the antiviral mechanism of a developmental host-directed therapeutic approach with broad applicability against emerging viral pathogens.

**Author Summary:** The COVID-19 pandemic has underscored the urgent need for effective countermeasures to novel and emerging viral pathogens. Our research presented here builds upon our previously published data on an experimental novel antiviral compound termed UMB18. We have found this compound capable of inhibiting replication of influenza A virus, coronaviruses and the filoviruses Marburg and Ebola virus, but did not fully define a mechanism of action. In this work, we demonstrate that UMB18 exerts antiviral activity by modulating cellular cholesterol levels. By targeting the SKI complex, UMB18 triggers an increase in endogenous cellular cholesterol which disrupts the fine balance viruses rely on for efficient infection. We demonstrate that this mechanism inhibits replication of SARS-CoV-2, revealing a previously undescribed host-directed strategy for antiviral intervention. These findings highlight UMB18’s potential as a broad-spectrum antiviral agent and pave the way for further research into its mechanism and therapeutic applications, offering a promising avenue for development of antiviral countermeasures to current, novel and emerging pathogens.

## Introduction

The coronavirus disease-2019 (COVID-19) pandemic caused by severe acute respiratory syndrome coronavirus-2 (SARS-CoV-2) has highlighted the lack of broadly acting therapeutics to treat coronaviruses and other emerging pathogens. These types of chemicals could serve as first-line countermeasures, filling a crucial gap before specific measures can be developed. Addressing this need, our research is focused on a host-directed approach of targeting the SKI (super killer) complex (1).

As opposed to direct acting antivirals, host-directed therapeutics have a significantly lower risk for the emergence of viral escape mutants, but such strategies come with a greater risk of side effects. It is therefore essential to understand the mechanism of action for any such approach.

Comprised of SKIC2, SKIC3, and SKIC8 (formerly SKIV2L, TTC37, and WDR61, respectively) the SKI complex acts as a cofactor to the cytosolic RNA exosome to facilitate 3’ to 5’ RNA degradation (2–4). SKIC2 is a member of the DExD/H helicase family, while SKIC3 and SKIC8 have structural and regulatory functions (5). The complex was first defined in yeast where it was implicated in resistance to the “super killer” toxin of a yeast double stranded RNA virus, hence the naming convention of SKI (6,7). Our previous study demonstrated that siRNA depletion of the SKI complex inhibited replication of influenza A virus (IAV) and Middle East respiratory syndrome coronavirus (MERS-CoV). We proceeded to use *in silico* modeling to predict chemicals that could hypothetically bind SKIC8 and potentially disrupt the proviral activity of the SKI complex. We demonstrated that our lead compound, termed UMB18, inhibited all of IAV, MERS-CoV, SARS-CoV, SARS-CoV-2, and the filoviruses Ebola virus and Marburg virus. Then, using single cycle IAV infections, we determined that UMB18 did not impact virus entry to cells, but potently inhibited RNA and subsequent protein production. Thus, we defined UMB18 as a host-directed antiviral, targeting post-entry steps prior to RNA production, with broad-spectrum activity (1).

In the present study, UMB18 was found to bind SKIC8 demonstrating specificity of our previously predicted binding. We subsequently found transcriptomic upregulation of genes involved in cholesterol synthesis and concomitant increases in total cholesterol following UMB18 treatment of cells. Cholesterol can be obtained from external sources through low-density lipoprotein (LDL) receptors or endogenously synthesized through the mevalonate pathway (8–10). Sterol regulatory element binding proteins (SREBPs) are master transcriptional regulators of cholesterol and fatty acid synthesis (11). SREBPs comprise three isoforms, SREBP1a, SREBP1c, and SREBP2, expressed from two genes *Srebf1* and *Srebf2* (12,13). Each of the SREBP proteins are regulated by SREBP-cleavage activating protein (SCAP) (14,15). We found that depletion of *Scap, Srebf1* or *Srebf2* reduced UMB18-mediated increases in mevalonate pathway genes. Notably, UMB18 antiviral activity against SARS-CoV-2 was significantly reduced with either depletion of the *Scap* or *Srebf* genes or extraction of cholesterol with methyl-β cyclodextrin. These results underscore the importance of increased total cellular cholesterol in the mechanism of action for UMB18 as an antiviral compound.

Overall, the work presented here demonstrates the promise of UMB18 as a broad-spectrum, host-directed antiviral compound and elucidates its mechanism of action through upregulation of cellular cholesterol synthesis by targeting the SKI complex.

## Results

### UMB18 binds SKIC8 and upregulates genes involved in cholesterol synthesis

We previously described the broad-spectrum antiviral activity of the SKI-targeted compound UMB18 (1) and now aimed to identify its mechanism of action. The compound was developed using *in silico* modeling and was predicted to bind to SKIC8 (formerly WDR61). The genes of the SKI complex are required for the antiviral activity of UMB18, suggesting specificity but we did not demonstrate that UMB18 directly binds to the predicted target SKIC8 (1). We therefore expressed and purified SKIC8 from bacterial cells and analyzed UMB18 binding by surface plasmon resonance (SPR). The fast association and dissociation rate constants observed at various compound concentrations indicated binding to the SKIC8 target with a relatively low binding affinity, KD = 81 μM (Fig. 1A). These data suggest that UMB18 is indeed directly binding to SKIC8.

**Figure 1.**
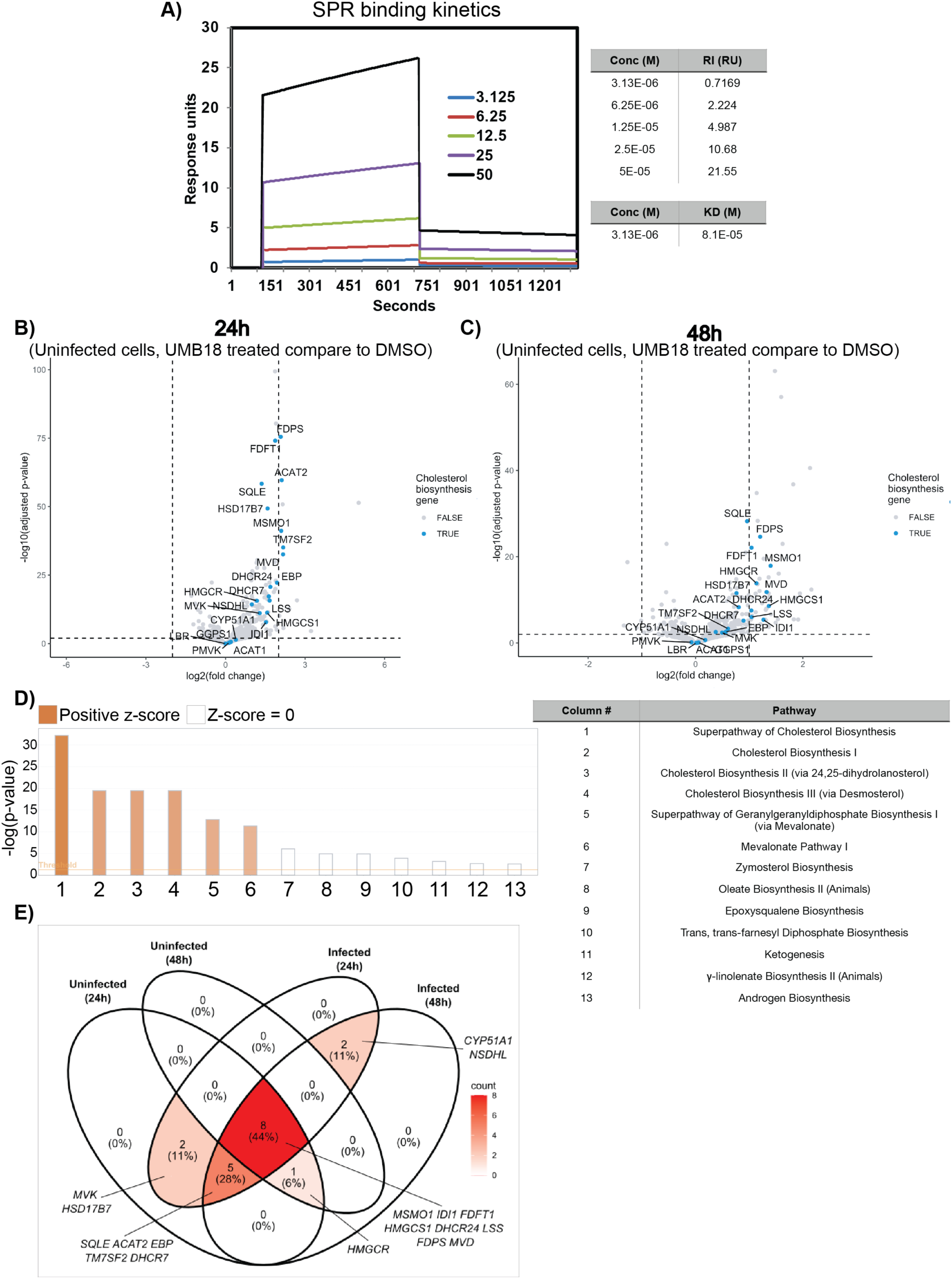
UMB18 directly binds SKIC8 and causes transcriptomic increases in mevalonate pathway. A) Binding of UMB18 to SKIC8. The binding affinity of immobilized SKIC8 to UMB18 at various concentrations was quantified by SPR as described in the methods. A KD of 81 μM was calculated for the binding. B) Volcano plots showing differential expression of genes at 24h (top) and 48h (bottom) in uninfected A549-ACE2 cells treated with 10 μM UMB18 compared to 0.1% DMSO treated, uninfected A549-ACE2 cells. Cholesterol biosynthesis genes are colored blue and labeled. A horizontal dashed line is drawn at *p_adj_* = 0.01, vertical dashed lines are drawn at +/- 2-fold change. C) Plot showing significantly enriched pathways in uninfected A549-ACE2 cells treated with UMB18 compared to DMSO treated, uninfected A549-ACE2 cells at 24h. Orange bars show pathways with predicted increased pathway output (z > 2); white bars show pathways with no predicted activity pattern (z = 0). D) Venn Diagram comparing upregulation of cholesterol biosynthesis genes in UMB18-treated A549-ACE2 cells against timepoint- and infection-matched, untreated controls. The number and color intensity in each area corresponds to the number of cholesterol biosynthesis genes that are significantly upregulated (*p_adj_* < 0.01) and these genes are indicated.

Having determined that our compound is specific to the *in silico* predicted target, we next utilized RNA transcriptomics to investigate the changes that UMB18 binding SKIC8 can cause, both in uninfected or SARS-CoV-2 infected cells. This provided an unbiased approach to investigate potential mechanisms of antiviral activity. A549 cells (human lung alveolar adenocarcinoma cells) overexpressing human angiotensin converting enzyme 2 (A549-ACE2) to allow for productive SARS-CoV-2 infection were used for the transcriptomic analysis and four conditions were analyzed: 1) 10 μM UMB18 treatment, 2) 0.1% DMSO treatment (carrier for UMB18), 3) 10 μM UMB18 with SARS-CoV-2 infection or 4) 0.1% DMSO with SARS-CoV-2 infection. We previously found 10 μM to be inhibitory to viral replication with no cytotoxicity (see (1) and also Fig. 6F). All infections were at multiplicity of infection (MOI) 0.1, performed across two experiments, with either 24-hour (h) and 48h or just a 24h of infection and treatment. In uninfected cells, the most prominently differentially expressed genes caused by UMB18 treatment compared to DMSO were in the mevalonate pathway, associated with cholesterol biosynthesis (Fig. 1B-D). As expected, SARS-CoV-2 infection had minimal impact on the genes that were upregulated with UMB18 treatment compared to DMSO, with the majority of significantly upregulated genes occurring in all conditions (Fig. 1E). Eight genes (MSMO1, IDI1, FDFT1, HMGCS1, DHCR24, LSS, FDPS, MVD) were significantly upregulated at both time points and both infection conditions, while five genes (SQLE, ACAT2, EBP, TM7SF2, DHCR7) were upregulated by UMB18 treatment at 24h and 48h infection as well as at the 24h but not the 48h time point in uninfected cells. MVK and HSD17B7 were upregulated at 24h in infected and uninfected cells, while CYP51A1 and NSDHL were upregulated in the context of infection at 24h and 48h but not in uninfected cells. HMGCR was upregulated at 24h and 48h in uninfected cells and at 48h in infected cells (Fig. 1E). These studies used bulk RNA sequencing with a low MOI of SARS-CoV-2 and 10 μM UMB18 treatment causes a roughly 90% inhibition of infection. As expected, low levels of SARS-CoV-2 replication are occurring in the treated cells (for example see Fig. 6F), therefore we did not expect to see major differences with infection. This data set was included for completeness and as a comparator to the uninfected cells. The analysis demonstrated upregulation of genes involved in the mevalonate pathway after treatment with UMB18 which may have a role in antiviral activity.

### UMB18 treatment increases cellular cholesterol levels

The upregulation of the mevalonate pathway through the transcriptomic analysis suggested that cholesterol synthesis may be affected by UMB18. We validated the results of the transcriptomic analysis using qRT-PCR with primers targeting a subset of the genes in the mevalonate pathway. Identical to the previous experiment, A549-ACE2 cells were treated with 10 μM UMB18 or 0.1% DMSO as the vehicle control for 24 or 48h. Additionally, A549 cells not expressing ACE2 were subject to the same treatments to further validate the results and ensure that ACE2 overexpression was not impacting the transcriptional changes. Seven genes for enzymes involved in various stages of the mevalonate pathway were analyzed to get a broad view of transcriptional changes (pathway schematic in Fig. 2A, yellow boxes are selected genes for analysis). Relative to DMSO treatment, UMB18 caused a significant increase in expression of genes in the mevalonate pathway, ranging from approximately 2- to 5-fold. In the A549-ACE2 cells (Fig. 2B), induction of the selected genes peaked at 24h and then declined by 48h. With 48h treatment, the genes were still more highly expressed than DMSO control cells but lacked the statistically significant differences seen at 24h (Fig. 2B). We suspect this may be a result of other feedback regulatory loops in the pathway or due to degradation of UMB18 in the media as it was not replenished through the time course. In A549 cells, UMB18 treatment caused an increase in expression of each of the genes that similarly peaked at 24h and declined at 48h. These changes remained statistically significant compared to DMSO controls at 48h (Fig. 2C). The magnitude of change between the two cell lines was largely similar. Overall, these qRT-PCR data suggest that UMB18 caused an increase in genes that regulate cholesterol synthesis through the mevalonate pathway and that the expression of ACE2, which allows for infection by SARS-CoV-2, does not significantly alter the magnitude of change.

**Figure 2.**
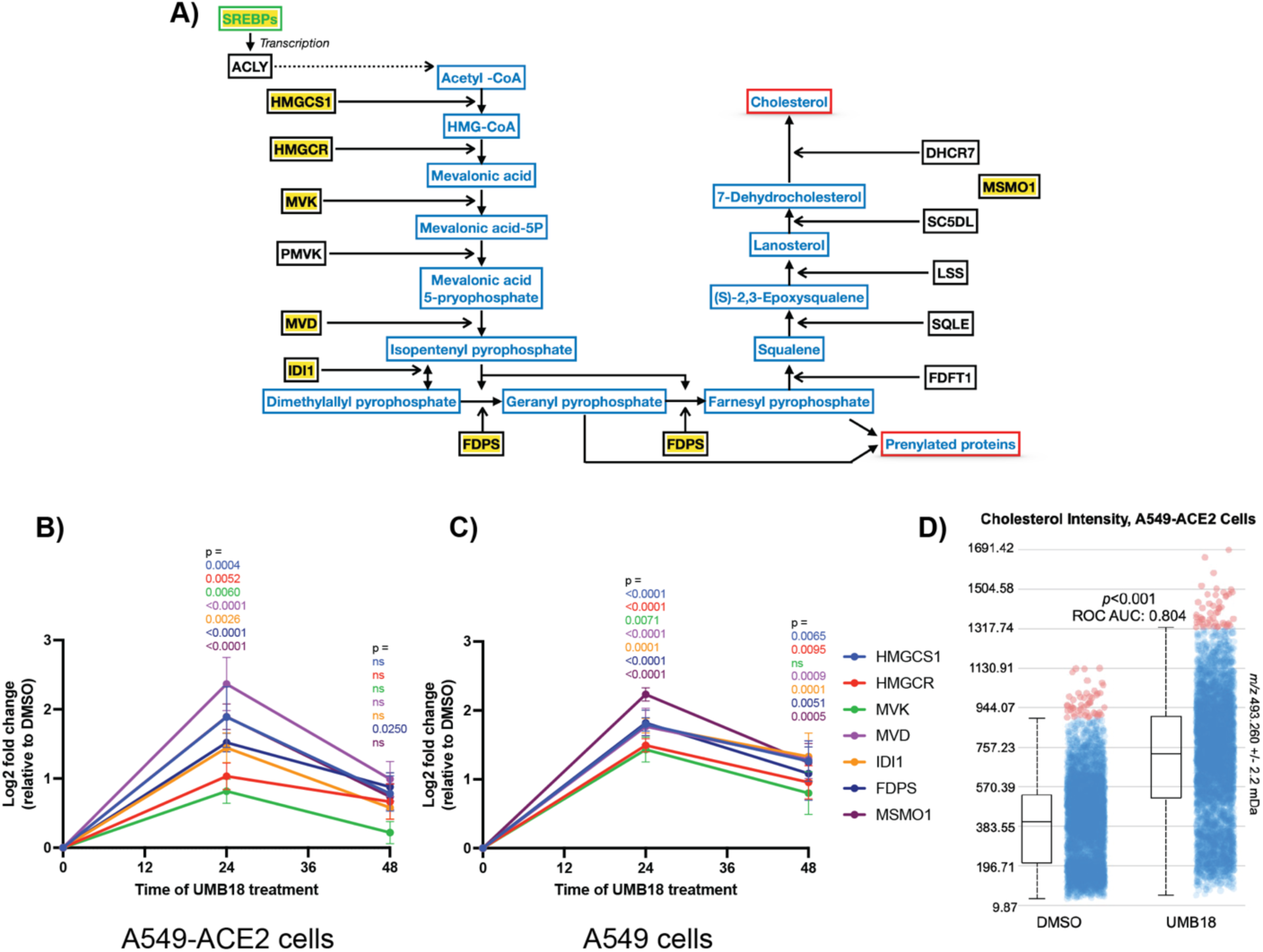
UMB18 causes increased expression of mevalonate pathway genes and cellular cholesterol. A) Simplified schematic representation of the mevalonate pathway leading to cholesterol synthesis. In blue are metabolic products of the pathway and in black are the enzymes that regulate the various steps with those that are highlighted being analyzed by qRT-PCR through the paper. B and C) A549-ACE2 cells (B) or A549 cells (C) were treated with 10 μM UMB18 or 0.1% DMSO vehicle control and collected at 24h or 48h in TRIzol for qRT-PCR analysis of expression of various genes throughout the mevalonate pathway. The Ct values were normalized to GAPDH as a housekeeping gene control to calculate the ΔCt and then relative fold change calculated between UMB18 treated cells and DMSO treated cells for the ΔΔCt. Plotted is the -ΔΔCt on a Log2 scale to show relative fold change, plotting mean and standard error of the mean as error bars. P values were calculated for the ΔCt values by two-way ANOVA with a Tukey multiple comparison correction for each gene at the different time points and treatments. Data are from 3 independent experiments each performed with triplicate wells of treatment. D) Relative intensity cloud projection with box plot of *m/z* 493.260 +/- 2.2 mDa ([cholesterol+Ag_107_]^+^) ion per image pixel for UMB18- and DMSO-treated cells with area-under-the-curve (AOC) from a receiver-operator characteristic analysis. Student’s T-test base on quadruplicate samples.

We next investigated whether total cellular cholesterol was increased in the cells, since we identified an increase in the expression of genes involved in the mevalonate pathway with UMB18 treatment. To this end, we utilized silver-assisted laser desorption ionization mass spectrometry imaging (Ag-LDI MSI), a highly sensitive method of detecting total cellular cholesterol. A549-ACE2 cell culture monolayers treated with UMB18 showed a higher intensity of cholesterol signal compared to DMSO-treated cells. Cholesterol was detected and visualized in the monolayers as the ion *m/z* 493.260 +/- 2.2 mDa ([cholesterol+Ag107]+) ion (Fig. 2D) and the silver isotopologue ion [cholesterol+Ag109]+ data agreed. Treatment with UMB18 resulted in a significant increase in relative cholesterol abundance, showing a nearly 2-fold increase in pixel intensities. Further, a receiver-operator characteristic analysis showed an area-under-the-curve value of 0.804 for cholesterol in UMB18 treated cells, making cholesterol abundance a strong discriminator over DMSO-treated cells. These data demonstrate that UMB18 treatment of A549-ACE2 cells results in significantly and discriminatorily higher amounts of cholesterol compared to control.

### UMB18 increases cellular cholesterol and exerts antiviral activity in serum containing media

Experiments to investigate changes caused by UMB18 treatment were performed in serum supplemented media and therefore have exogenous cholesterol available. Cells can endocytose cholesterol through the LDL-receptor and activate a negative feedback loop to inhibit the mevalonate pathway. It therefore appeared UMB18 may activate this metabolic pathway even in the presence of exogenous cholesterol. To further analyze the system, we treated A549-ACE2 cells with 10 μM UMB18 or 0.1% DMSO in serum-containing or serum-free media, isolated RNA at 24h, and performed qRT-PCR for a subset of the mevalonate pathway genes. In serum-containing media, UMB18 treatment resulted in a similar significantly increased gene expression for the mevalonate pathway genes HMGCS1, MVD and MSMO1 (Fig. 3A-C). In serum-free media, there was an upregulation of these three genes, in both DMSO and UMB18 containing media, with no significant differences between the two. Serum-free media caused a statistically significantly larger increase in expression compared to UMB18 in serum-containing media in all cases (Fig. 3A-C). These data show that in the absence of exogenous cholesterol from serum, the mevalonate pathway was activated as expected and, in that context, UMB18 did not further enhance the level of mevalonate pathway associated gene transcription. However, in media containing serum, UMB18 did cause a statistically significant increase in genes involved in the mevalonate pathway.

**Figure 3.**
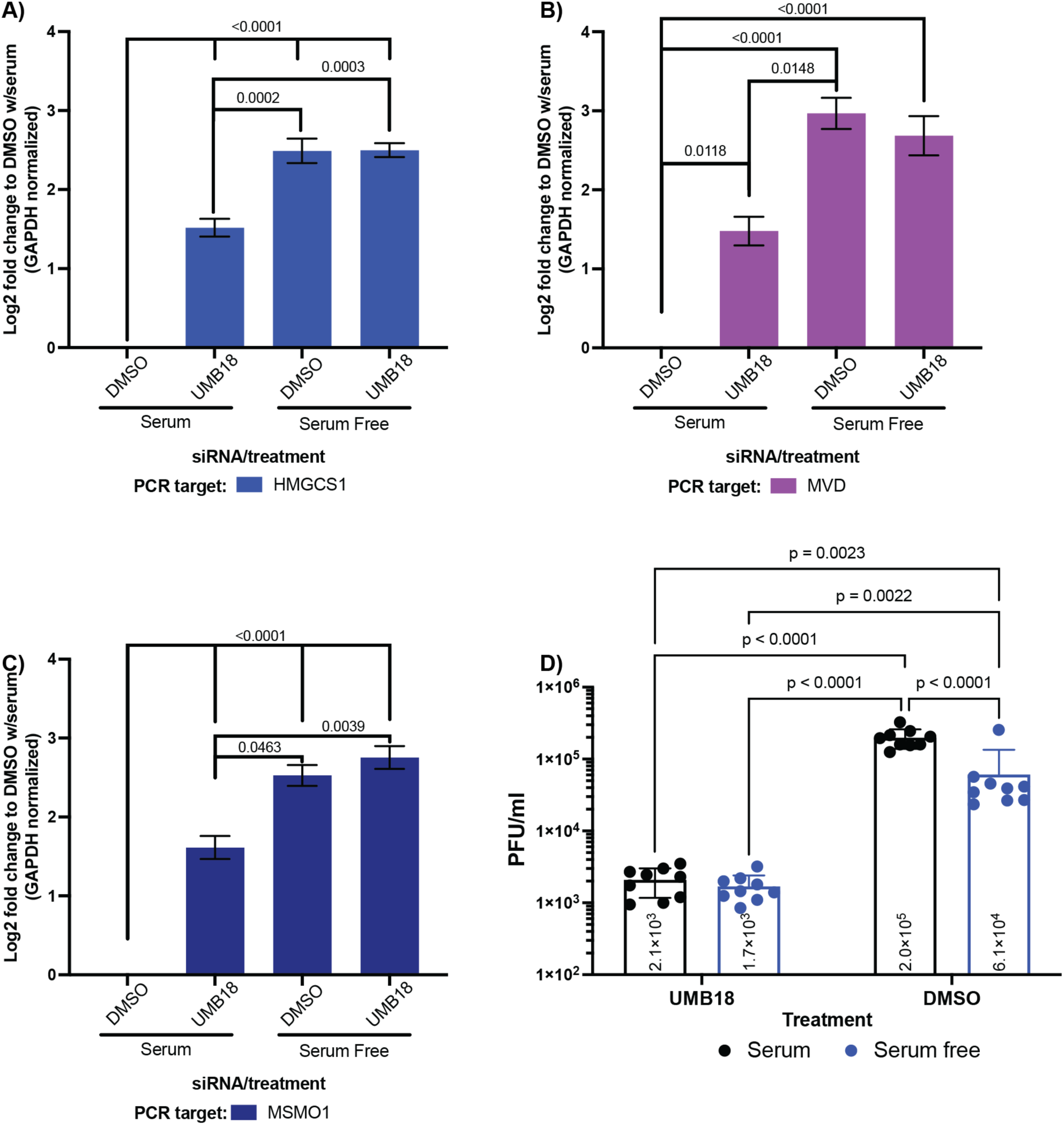
UMB18 increases cellular cholesterol and exerts antiviral activity in serum containing media. A-C) A549-ACE2 cells were treated with 10 μM UMB18 or 0.1% DMSO in culture media containing 10% (v/v) fetal bovine serum (as described in Materials & Methods) or without supplemental serum (serum free). After 24h of treatment, cells were collected in TRIzol for qRT-PCR analysis of the three indicated genes (HMGCS1, MVD and MSMO1, A-C). Analysis was performed, and data plotted as described in Figure 2. D) A549-ACE2 cells were infected with SARS-CoV-2 at MOI 0.5 in serum containing or serum free media with 10 μM UMB18 or 0.1% DMSO. 24-hour post-infection supernatant was collected and used to titer virus by plaque assay on VeroTMPRSS2 cells. Plotted is the mean pfu/ml value from 3 independent experiments performed in triplicate wells, with that value being displayed within each bar. Significance calculated by two-way ANOVA with a Tukey multiple comparison correction. Only statically significant (p < 0.05) comparisons are shown on the graph.

We hypothesized that this increase in cholesterol synthesis caused by UMB18 plays a role in antiviral activity against SARS-CoV-2. We therefore infected A549-ACE2 cells with SARS-CoV-2 at MOI 0.1 in serum-containing or serum-free media and treated with 10 μM UMB18 or 0.1% DMSO for 24h prior to supernatant collection and quantitation of virus by plaque assay. UMB18 was equally effective at inhibiting viral replication regardless of the presence of serum (Fig. 3D). In serum-free media containing DMSO, there is a small (3x), but statistically significant drop in viral titer compared to serum-containing media, suggesting that the increased expression of mevalonate pathway genes may have an impact on viral replication (Fig. 3D). The inhibition is not as potent as UMB18 treatment, and we hypothesize this may be a temporal effect with UMB18 causing a more rapid increase in expression of the genes that have activity and inhibit early stages of the viral life cycle.

### SCAP and SREBP are required for UMB18-induced increases in mevalonate pathway genes

Our data suggest that UMB18 binds to the SKI complex and consequently increases expression of genes in the mevalonate pathway leading to an increase in total cellular cholesterol. We set out to determine what cellular pathways the SKI complex might be interacting with to mediate this effect. First, we looked at the mammalian target of rapamycin (mTOR) complex because of published findings that demonstrated a loss of SKIC2 activity was connected to an increase in mTORC1 signaling (16). The mTOR complex is a master regulator of numerous aspects of cellular metabolism and it can regulate activation of the sterol regulatory element binding proteins (SREBPs) which promote expression of the mevalonate genes. We analyzed the ribosomal protein S6 kinase 1 (S6K1) which is phosphorylated when the mTOR pathway is activated. Using treatment with epidermal growth factor (EGF), a known activator of the mTOR pathway, we were able to detect phospho-S6K1 (pS6K1), demonstrating that our cells showed a canonical marker for mTOR signaling (Fig. 4A). We proceeded to treat cells with UMB18 for various time points for analysis (Fig. 4B-C). None of these treatments showed any significant changes to phospho-S6K1, suggesting that UMB18 did not cause a detectable activation of mTOR, at least based on phosphorylation of S6K1.

**Figure 4.**
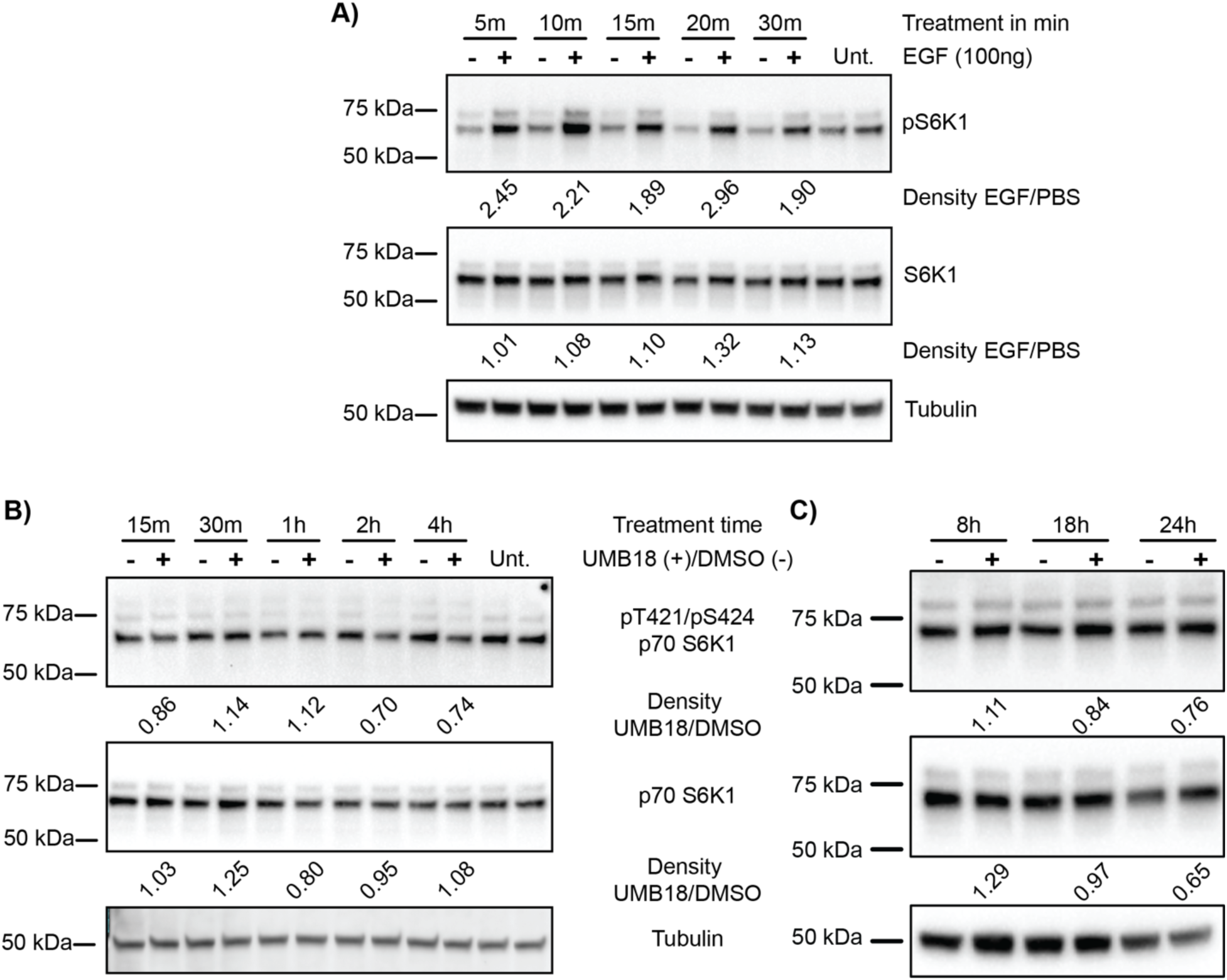
UMB18 does not appear to activate mTOR signaling as assessed by phospho-S6K1. A) A549-ACE2 cells were plated to plastic and the following day treated with 100 ng recombinant EGF, or PBS carrier alone. At time points between 5 and 30 min, cells were collected in Triton-X100 lysis buffer and prepared for western blotting (see Materials and Methods). Samples were separated on gels and blotted for phospho-S6K1 or S6K1 and tubulin was used as a loading control. All probes were detected with HRP. Density of bands was determined and quantified by setting relative to tubulin control and then reported as a relative band density of EGF treated compared to PBS treated at each time point. B and C) A549-ACE2 cell were plated to plastic and the following day treated with 10 μM UMB18 or 0.1% DMSO for the indicated time points or left untreated (for 4 hours in B). At each timepoint, cells were collected, and western blots run similarly to A. All gels were imaged with HRP, except tubulin in B which was visualized by AlexaFluor 488. Density of UMB18 compared to DMSO at each time point is reported as in A.

Since UMB18 did not show any clear activation of mTOR, we next investigated the master regulators of cholesterol and fatty acid synthesis, the SREBPs. Two genes, *Srebf1* and *Srebf2,* produce three proteins, SREBP-1a, SREBP-1c, and SREBP2 and all of these are regulated by SREBP cleavage activating protein (SCAP), encoded by the *Scap* gene (11,13,14,17). Transfection of siRNAs targeting each of these three genes into A549-ACE2 cells was used for knockdown studies. At 72h post siRNA transfection, cells were treated with DMSO or UMB18 for 24h and RNA collected to perform qRT-PCR for each of the siRNA targets and three mevalonate pathway genes (Fig. 5). We hypothesized that if UMB18 was activating mevalonate pathway gene expression through the canonical route, reduced expression of SCAP or SREBPs would reduce the impact of this treatment.

**Figure 5.**
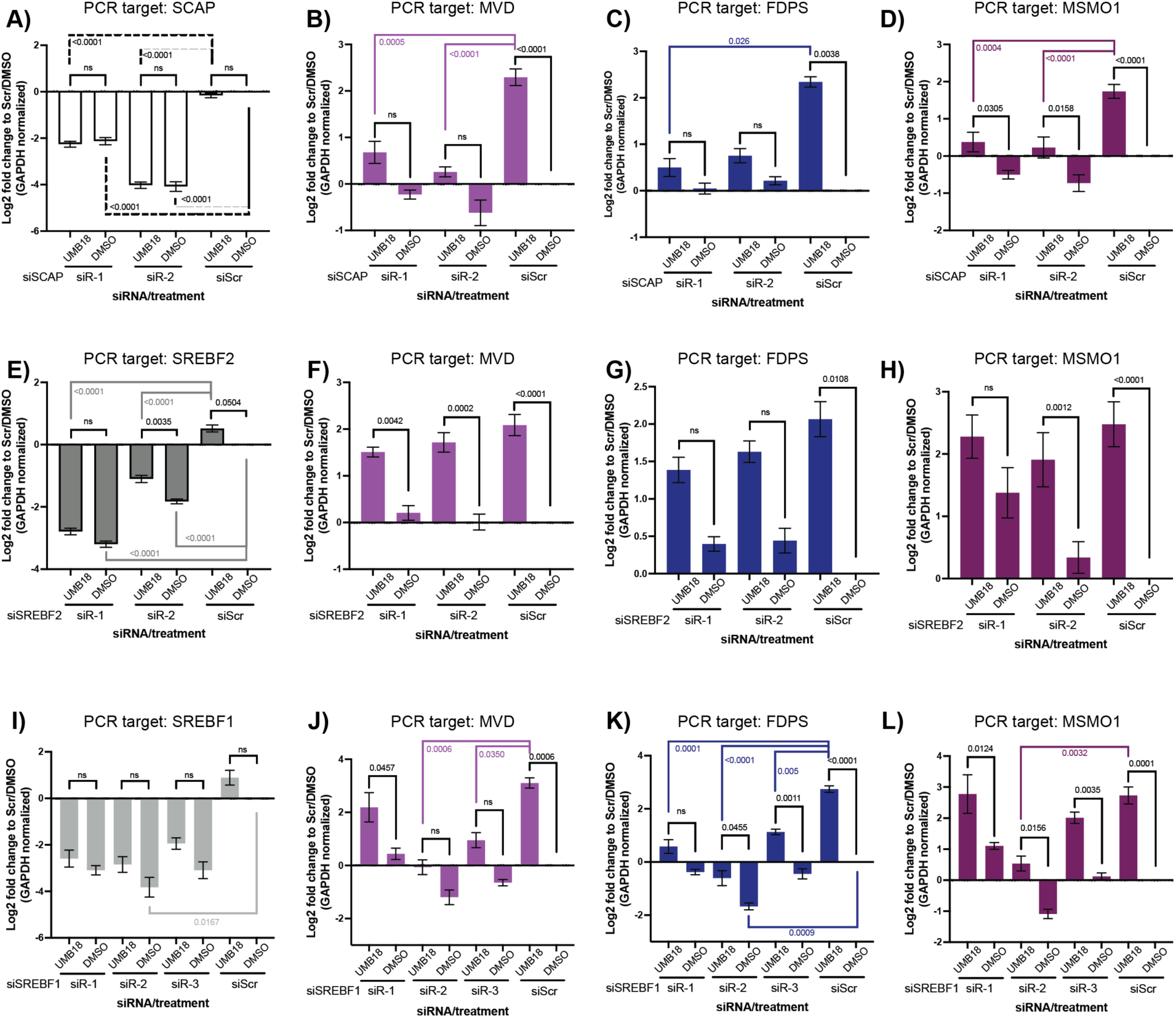
UMB18 requires SCAP and SREBP genes to increase mevalonate pathway gene expression. A549-ACE2 cells were transfected with unique siRNA sequences targeting *Scap* (A-D)*, Srebf2* (E-H) or *Srebf1* (I-L) for 72h. The cells were then treated with 10 μM UMB18 or 0.1% DMSO for 24h and collected in TRIzol for qRT-PCR targeting the specified genes (the siRNA targeted genes or MVD, FDPS or MSMO1 as representatives of the mevalonate pathway). The Ct values were normalized to GAPDH as a housekeeping gene to determine the ΔCt and then relative fold change was calculated compared to scrambled siRNA (siScr) transfected cells treated with DMSO (siScr/DMSO) for each of the conditions for the ΔΔCt. The mean -ΔΔCt is plotted on a Log2 scale to show relative fold change, plotting mean and standard error of the mean as error bars. P values were calculated for the ΔCt values by two-way ANOVA with a Tukey multiple comparison correction for each gene. Values were calculated comparing the mean within each siRNA condition (UMB18 vs DMSO), displayed with black bars or within each treatment condition where gene specific knockdown siRNA was compared to siScr for UMB18 treatment (color bars above graph) or DMSO treatment (color bars below graph). For this later comparison, only statistically significant p values (< 0.05) are displayed. Data are from 3 independent experiments each performed with triplicate wells of treatment.

SCAP is the master regulator for activation of the SREBPs. We targeted SCAP with two different siRNA sequences and both caused a significantly decreased expression of *Scap* relative to scrambled siRNA (siScr), with sequence 2 being more potent that sequence 1 (Fig. 5A). There was no significant impact of UMB18 treatment upon expression of *Scap* in any context (Fig. 5A). In siScr transfected cells treated with UMB18, there was a significant increase in expression of *MVD*, *FDPS* and *MSMO1* compared to DMSO treated cells (Fig. 5B-D, right most pairs). Each gene had 4 times the expression level based on the qRT-PCR data, matching previous results (Fig. 2 and 3). In the context of *Scap* siRNA there was no significant difference between UMB18 and DMSO treated cells for *MVD* and *FDPS* gene expression (Fig. 5B and C), but there were significant changes for *MSMO1* (p = 0.0305 and 0.0158 for siRNA sequence 1 and 2; Fig 5D). However, the relative fold change was significantly lower than that caused by UMB18 in siScr transfected cells (both p < 0.0001). These data all suggest that UMB18 did not impact the expression of *Scap*, but when expression of this gene was reduced through siRNA-mediated knockdown there was a significantly reduced impact of UMB18 on the mevalonate pathway genes. This suggests that UMB18 treatment activates expression of these genes through a canonical pathway mediated by SCAP.

To further validate that UMB18 is impacting this canonical signaling pathway, we also targeted the *Srebf* genes with siRNA. *Srebf2* was knocked down with 2 separate siRNA sequences similarly to *Scap* and both caused a significant decrease in expression. Sequence 1 was more potent than 2, however both caused a significant decrease in gene expression (Fig. 5E). UMB18 did have some impact on *Srebf2*, unlike the results for *Scap*, but that could be attributed to the increase in mevalonate pathway genes since *Srebf2* is in a positive feedback loop. In the context of siScr there was a mild increase in expression, and the p value was calculated to 0.0504. Similarly, in the context of siRNA sequence 2, there was significantly less reduction in *Srebf2* expression cause by UMB18 treatment, with a p value of 0.0035 between UMB18 and DMSO conditions. However, compared to siScr transfection and DMSO treatment, the UMB18 treated cells still had an approximate 2-fold decrease in expression, suggesting any impact from UMB18 does not outweigh the inhibition from the siRNA. For each of the mevalonate pathway genes, once again, UMB18 caused a significant increase in expression compared to DMSO within the control condition of siScr transfection (Fig 5F-H, right most pairs). Loss of *Srebf2* appeared to have less impact on the UMB18-mediated changes than *Scap* with significant changes observed in both siRNA transfection conditions for *MVD* (Fig 5F), and the context of siRNA sequence 2 for *MSMO1* (Fig. 5H). However, the changes in *FDPS* were not significant between UMB18 and DMSO with either siRNA sequence (Fig 5H). and for sequence 1 for *MSMO1* expression (Fig 5H, left most pair). These data suggest that reduced *Srebf2* expression through siRNA reduced the impact UMB18 had on altering expression of these three mevalonate pathway genes, however, the targeting of *Scap* with siRNA appeared to cause a more potent inhibition.

Finally, to complete this investigation of the SCAP/SREBP pathway, *Srebf1* was targeted with siRNA, with 3 different siRNA sequences. While all of the sequences caused approximately a 4x or greater decrease in expression, these data did not reach statistical significance, with the exception of siRNA sequence 2 compared to siScr in the context of DMSO treatment (Fig 5I). UMB18 did not significantly impact expression of *Srebf1* (Fig. 5I). As for other experiments, UMB18, in the context of siScr transfection, caused a significant increase in expression of *MVD*, *FDPS* and *MSMO1* compared to DMSO treatment (Fig. 5J-L, right most pairs). Even though the three siRNA sequences did not consistently cause a statistically significant decrease in expression, all of them showed varying levels of decrease for UMB18 impact on gene expression. For *MVD*, siRNA sequence 2 and 3 both had a significantly reduced fold change for UMB18 treatment (Fig 5J, p = 0.0006 and 0.0350). For *FDPS*, all three sequences had significantly reduced fold changes (Fig. 5K) and for siRNA sequence 2 there was a significant reduction in UMB18 induced changes to *MSMO1* expression (Fig. 5L). These data suggest that even though these siRNA sequences did not cause statistically significant reduction in *Srebf1* expression, all of them caused significant inhibition of UMB18 induced changes.

Overall, the data in Fig. 5 demonstrate that UMB18 caused an increase in expression of genes in the mevalonate pathway (represented by *MVD*, *FDPS* and *MSMO1*) without causing major changes to *Scap*, nor the *Srebf* genes. However, when the SCAP/SREBP axis is disrupted by siRNA treatment, there were significant reductions to the impact from UMB18 on expression of mevalonate pathway genes. Our results demonstrate that UMB18 directly binds to the SKI complex (Fig. 1A) and causes genetic activation of the mevalonate pathway (Fig. 2, 3 and 5) which relies on the canonical activation of SCAP/SREBPs.

### Increased cellular cholesterol is required for UMB18-mediated antiviral activity

We have found that UMB18 caused an increase in mevalonate pathway gene expression and that the canonical signaling pathway mediated by SCAP and SREBPs is involved. This transcriptional response increased total cellular cholesterol, leading to the question of whether the increase in cholesterol is important for antiviral activity. To investigate this, we again used siRNA to target *Scap* and the *Srebf* genes and treated cells with UMB18 while also infecting with SARS-CoV-2. A549-ACE2 cells were plated and transfected for 72h. Cells were then infected with SARS-CoV-2 at MOI 0.1 and treated with 10 μM UMB18 or 0.1% DMSO for 24h. Supernatant were collected, and viral titer determined by plaque assay on VeroE6 cells expressing transmembrane protease serine 2 (TMPRSS2). With scrambled siRNA there was an approximate 10-fold drop in viral titer (Fig. 6A-C), similar to the effects we previously published (1). However, when *Scap* was knocked down by siRNA, UMB18 lowered viral titer by 4.36- and 2.92-fold compared to DMSO and was not significantly different (Fig. 6A and C). These data suggest the reduction in *Scap* expression caused by siRNA reduced the antiviral activity of UMB18, making it not statistically significant compared to DMSO. When *Srebf2* was targeted by siRNA sequence 1 there was only a 2.05-fold difference between UMB18 and DMSO (but this was statistically significant), while sequence 2 had less impact with a 7.52-fold difference. These differences are in alignment with the potency of the siRNA sequences for knockdown of *Srebf2*, with sequence 2 being less potent (Fig. 5E) and causing less impact to UMB18 antiviral activity. For Srebf*1* knockdown down, UMB18 lowered viral titer by only between 1.23- to 2.04-fold, compared to DMSO (Fig. 6C), suggested even great loss of inhibition. In all cases, viral titer inhibition due to UMB18 is attenuated by loss of SCAP and SREBPs, with varying degrees of significance. Since the loss of these genes causes a decrease in mevalonate pathway gene expression induced by UMB18 (Fig. 5), these data collectively suggest that increased cellular cholesterol plays a role in UMB18-mediated antiviral activity.

**Figure 6.**
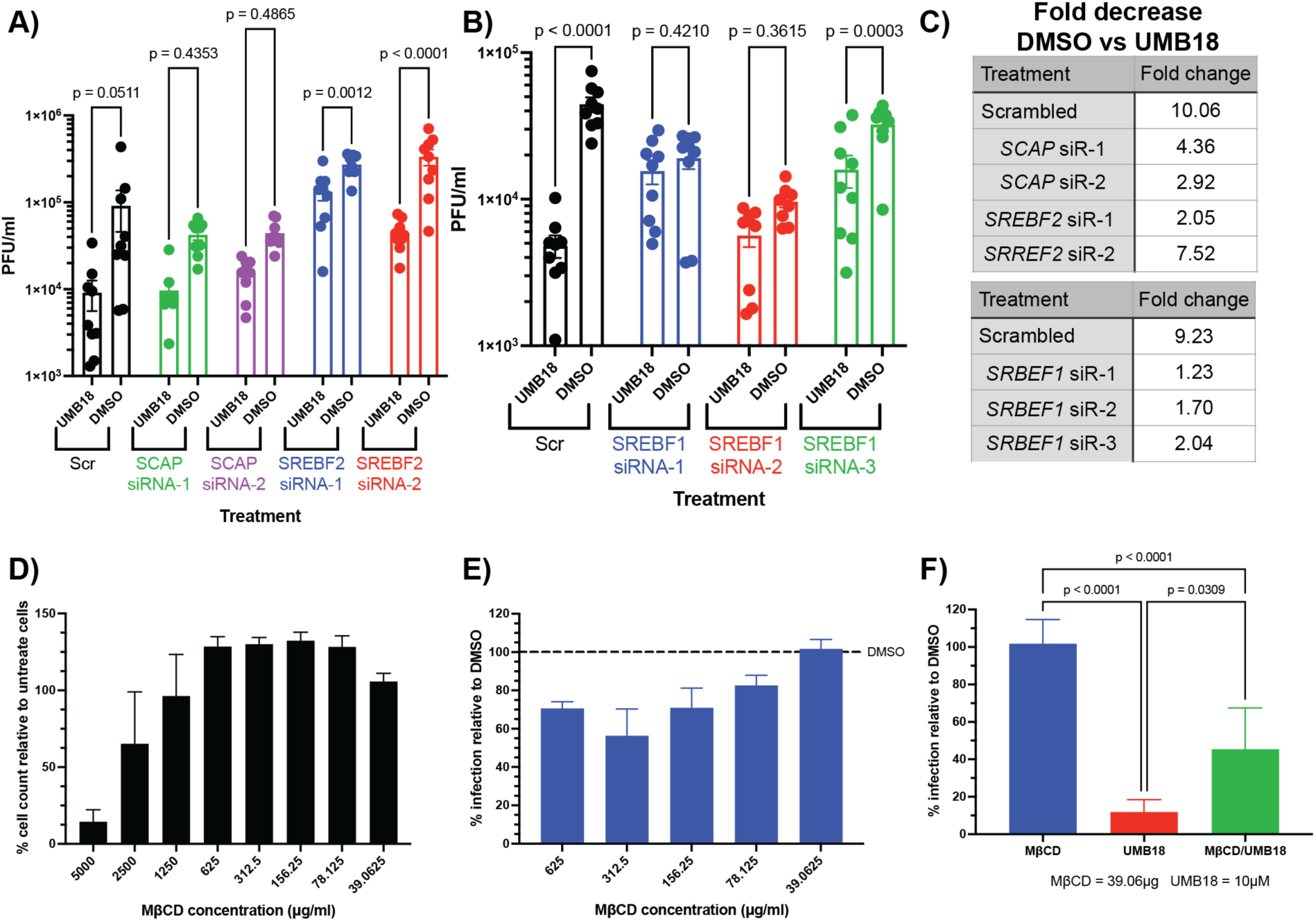
Increased cellular cholesterol is required for UMB18-mediated antiviral activity. A and B) A549-ACE2 cells were transfected with siRNA sequences targeting *Scap or Srebf2 or Srebf1* (sequences as used in Figure 5, *Scap* and *Srebf2* [A] transfections were experimentally performed together while *Srebf1* [B] were performed separately). Following transfection, cells were infected with SARS-CoV-2 at MOI 0.1 and treated with 10 μM UMB18 or 0.1% DMSO for 24h. Supernatant was collected from the cells and viral titer determined by plaque assay on VeroTMPRSS2 cells. The titer is plotted as plaque forming units/ml from each of the different transfection and treatment conditions. Using scrambled siRNA (siScr) as the control. Two-way ANOVA analysis was performed, and the p-values are plotted. Data are from three independent experiments performed in triplicate. C) Fold change between titer of DMSO treated and UMB18 treated in each transfection condition summarized in a table format. D) A549-ACE2 cells were treated with the indicated concentrations of methyl-beta-cyclodextrin (MβCD) for 24h prior to fixation with neutral buffered formalin. Cells were stained with Hoechst to label nuclei and cell count per well (96 well format) was quantified with a Celigo high content imager. Reported is the percentage total cell count compared to untreated cells as a measure of cytotoxicity from MβCD treatment. E) A549-ACE2 cells were infected with SARS-CoV-2 at MOI 3 and treated with the indicated concentrations of MβCD for 24h. Cells were similarly fixed with neutral buffered formalin then immunofluorescence labeled for SARS-CoV-2 N protein and stained with Hoechst. High content imaging was used to quantify percentage infection, and the plotted data are relative parentage infection compared to untreated control cells. F) As in E, cells were infected with SARS-CoV-2 at MOI 3 and treated with or without MβCD with either 10 μM UMB18 or 0.1% DMSO. Cells were similarly fixed and labeled for high content imaging of percentage infection. Plotted are the relative infection percentage to 0.1% DMSO treated cells for each of the different conditions. An unpaired t-test was performed for each condition and the p-values are reported. Data are from three independent experiments.

We also investigated the effects of methyl-β cyclodextrin (MβCD), a chemical that extracts cholesterol from cells, to see if this would disrupt UMB18 antiviral activity. Initially, A549-ACE2 cells were treated with a range of concentrations between 39 and 5000 μg/ml to investigate toxicity. The higher concentrations resulted in cell death based on automated counting of nuclei in a 96-well plate format (Fig. 6D). At 625 μg/ml and lower, there were no signs of toxicity. A549-ACE2 cells were then infected with SARS-CoV-2 at MOI 3 and treated with MβCD at non-cytotoxic concentrations. At 24 hours post-infection cells were fixed and immunofluorescent labeled for the viral N protein and high content imaged to determine the percentage infection. Fig. 6E demonstrates that MβCD has some level of antiviral activity with 625, 312.5, and 156.25 μg/ml causing roughly 30-40% fewer infected cells compared to DMSO-treated cells. However, at 39 μg/ml, there was no antiviral activity from MβCD alone. We, therefore, combined 39 μg/ml MβCD with 10 μM UMB18 to investigate whether extracting cholesterol from cells treated with UMB18 would reduce the antiviral activity. Fig. 6F demonstrates that UMB18 is a potent inhibitor, reducing percentage infection by about 90% compared to DMSO-treated control cells. When combined with MβCD, the percentage inhibition is approximately 40%, demonstrating that MβCD significantly reduced the antiviral activity of UMB18. These data demonstrate that reduced expression of the *Scap* or *Srebf* genes or cholesterol extraction from cells significantly reduced the antiviral activity of UMB18, pointing towards increased cholesterol being the antiviral mechanism of action for UMB18.

## Discussion

We have identified the SKI complex as a potential host-directed antiviral drug target (1) and determined that the novel compound, UMB18, specifically binds to its *in silico* predicted target, SKIC8. RNA transcriptomics, validated by qRT-PCR, demonstrated that UMB18 treatment caused significant upregulation of genes involved in the mevalonate pathway, and this corresponded to an increase in total cellular cholesterol (Fig. 1 and 2). Interestingly, these results were in the context of media supplemented with fetal bovine serum which contains cholesterol that can inhibit activation of the mevalonate pathway through a negative feedback loop. We found that cells cultured in serum-free media activated the mevalonate pathway and UMB18 had minimal additive effect (Fig. 3). We propose that UMB18 cannot further increase signaling to the already active pathway. However, we cannot rule out alternative hypotheses that UMB18 may be modified by components of serum, may not be effectively taken into cells in the absence of serum or may have impacts on roles adjacent to the mevalonate pathway such as protein prenylation. We do suspect some of these may have a role since virus replication is attenuated in serum-free media but UMB18 still provides a greater level of inhibition (Fig. 3D). As this compound is further developed, biochemical experiments to look at the impact of serum on its activity will be performed.

With the observed increase in expression of mevalonate pathway genes we were drawn to the SCAP/SREBP axis as the master regulators of sterol and fatty acid synthesis (11). Depletion of any of *Scap, Srebf1*, or *Srebf2* by siRNA resulted in an attenuation of UMB18-induced upregulation of mevalonate pathway genes, indicating that the UMB18 effect is through this canonical signaling pathway (Fig. 5). Crucially, the antiviral activity of UMB18 against SARS-CoV-2 was markedly decreased when *Scap* or *Srebf* genes were depleted from cells, arguing that the increased expression of mevalonate pathway genes is essential for antiviral activity (Fig. 6). Consistently, extraction of cellular cholesterol using MβCD also attenuated the antiviral activity of UMB18, reinforcing the importance of increased cellular cholesterol in mediating antiviral activity (Fig. 6).

Our findings collectively reveal that UMB18 binds the SKI complex and activates SREBP-mediated gene expression, leading to an increase in cellular cholesterol which is crucial for antiviral efficacy of the compound. We have shown that UMB18 and similar SKI-targeting compounds exhibit broad-spectrum activity against enveloped viruses, including coronaviruses, orthomyxoviruses, and filoviruses (1), all of which rely on manipulation of cellular membranes for efficient replication. We hypothesize that UMB18-mediated increase in total cellular cholesterol, which causes an increase in membrane rigidity, impedes the ability of these viruses to manipulate membranes and thus disrupts efficient replication.

Previous studies have suggested SARS-CoV-2 infection may downregulate expression of mevalonate pathway genes associated with cholesterol synthesis (18,19). It would therefore be reasonable to hypothesize that increasing expression of these genes would have antiviral activity, in alignment with our data. However, contrasting studies have suggested that SARS-CoV-2 and other coronaviruses cause an increase in lipid metabolism, mediated by the SREBPs (20–22), and that lowered cellular cholesterol can be antiviral (23,24). We also present data from our MβCD studies that lowered cellular cholesterol can be antiviral (Fig. 6E). Therefore, we propose that there may be a delicate equilibrium in cholesterol levels, or a “Goldilocks” range of cellular cholesterol necessary to support viral replication. Deviations from this balance, whether through insufficient or excessive cholesterol, disrupts membrane dynamics and adversely affects viral replication. At an organismal level, increased cholesterol usually comes from, or with, various other underlying health conditions which can all impact viral disease pathogenesis. In future work, we intend to further study the role these cellular cholesterol changes have at an organismal level using animal models of UMB18 treatment.

One hypothesis for the mechanism through which UMB18 is causing the SREBP-mediated transcriptional changes draws from work by Yang *et al.* (16), which linked SKIC2 deletion in mice to an increase in mTORC1 signaling. Despite efforts to investigate whether UMB18 activated mTORC1 as assessed by phosphorylation levels of S6K1, we observed no significant changes (Fig. 4). The activation of mTORC1 drives many cellular changes, one of which is the activation of the SREBP proteins, and thus a link between the SKI complex and mTORC1 activation could make sense in the context of our data. It is possible that UMB18 could activate mTORC1 but not at a level to observe changes to the phosphorylation status of S6K1.

Alternatively, UMB18 may alter the activity the RNA helicase activity of the SKI complex and impact genes that regulate SCAP/SREBP proteins independently of mTORC1. For instance, the SKI complex has been shown to have a role in resolution of stalled ribosomes (25) which we hypothesize to be disrupted by treatment with UMB18. Stalled ribosomes can activate ZAKα, which can in turn activate JNK signaling (26) and JNK signaling has been linked to activation of SREBPs (27). Future research will explore the interaction between UMB18, the SKI complex and these various cellular signaling pathways.

This study advances our understanding of our developmental host-directed, broad-spectrum antiviral therapeutic targeting the SKI complex. Building on our previous work which established the SKI complex as a host-directed target to inhibit diverse highly pathogenic human viruses, we reveal a crucial role for increased cellular cholesterol in mediating this antiviral effect. Our findings prompt further investigation into the mechanisms by which the SKI complex, an RNA helicase associated with 3’ to 5’ RNA degradation, influences metabolic pathways crucial for lipid and cholesterol synthesis, a finding that will be the focus of future work.

## Material and Methods

### Mammalian cell culture

A549 cells and A549 cells overexpressing human ACE2 (A549-ACE2, kindly provided by Dr. Brad Rosenberg (MSSM, NYC)) were cultured in DMEM (Quality Biologicals), supplemented with 10% (v/v) heat inactivated fetal bovine serum (FBS; Sigma) and 1% (v/v) penicillin/streptomycin (pen/strep, 10,000 U/ml / 10 mg/ml; Gemini Bio-Products). VeroE6 cells overexpressing TMPRSS2 (VeroT, kindly provided by Shutoku Matsuyama (National Institute of Infectious Diseases, Japan - Shirato2018_Virology) were cultured in DMEM, supplemented with 10% FBS, 1% pen/strep and 1% (v/v) L-glutamine (2 mM final concentration, Gibco). Cells were maintained at 37°C and 5% CO_2_.

### Chemical compounds

SKI targeting compounds were synthesized by, and purchased from, Dalriada Drug Discovery and were previously described (1). Methyl-beta-cyclodextrin (MβCD) was purchased from Sigma.

### Viruses and infections

Details on stock production have been described previously (1). SARS-CoV-2 strain Washington 1 (WA1) was initially provided by the CDC (BEI #NR-52281). All coronavirus work was performed in a Biosafety Level 3 laboratory and approved by our Institutional Biosafety Committee. In all cases of infection for experiments, cells were plated one day prior to infection to be at a desired confluence for infection. All viral titer were determined using plaque assays on VeroT cells as described (28). For experimental infections, virus was diluted to indicated MOI in media used for culture of A549 cells with addition of chemical compounds or controls as appropriate. For experiments using MβCD, 50 mM Hepes (Gibco) was additionally added to the media.

### RNA extraction and qRT-PCR

RNA extractions and cDNA reactions were performed as described previously (1) using Direct-zol RNA miniprep kit (Zymo Research) and RevertAid RT Kit (Thermo Scientific). The primers used are listed in the table below. A QuantStudio 5 (Applied Biosystems) was used for the qRT-PCR reactions. Loading was normalized using GAPDH and fold change was determined by calculating ΔΔCT after normalization.

#### RT-PCR primers

All primers for qRT-PCR were purchased from Integrated DNA Technologies (IDT) using their pre-designed qPCR assay primers:

| Gene target | IDT Assay ID |
| --- | --- |
| **SKIC2 (A.K.A SKIV2L)** | Hs.PT.58.14722008 |
| **SKIC3 (A.K.A TTC37)** | Hs.PT.58.26762302 |
| **SKIC8 (A.K.A WDR61)** | Hs.PT.58.157001 |
| **HMGCS1** | Hs.PT.58.4084870 |
| **HMGCR** | Hs.PT.58.41105491 |
| **MVK** | Hs.PT.58.40379926 |
| **MVD** | Hs.PT.58.39035173 |
| **IDI1** | Hs.PT.58.1930362 |
| **FDPS** | Hs.PT.58.27570524 |
| **MSMO1** | Hs.PT.58.3747602 |
| **SCAP** | Hs.PT.58.45442299 |
| **SREBF1** | Hs.PT.58.3359761 |
| **SREBF2** | Hs.PT.58.45335433 |
| **GAPDH** | Hs.PT.39a.22214836 |

### Transcriptomic sequencing and analysis

Library preparation and sequencing were carried out by the University of Maryland Institute of Genomic Sciences (IGS; Baltimore, MD, USA) on an Illumina NovaSeq 6000; 100bp paired-end sequencing on an S4 flow cell (Illumina, San Diego, CA, USA). Reads were preprocessed using cutadapt v3.4(29), then aligned to the *Homo sapiens* genome with STAR v2.7.8(30). Gene expression analysis was carried out using DESeq2 v4.1.0(31) in R (Rstudio, Boston, MA, USA). Pathway analysis was carried out using Ingenuity Pathway Analysis (QIAGEN, Hilden, Germany) using an alpha value of *p_adj_* < 0.01 to call a gene significantly differentially expressed. Raw sequence data is available in the NCBI Sequence Read Archive under the accession number PRJNA1055076.

### Mass spectrometry imaging

A modified cell culture apparatus was used for compatibility with mass spectrometry imaging (MSI). Briefly, Superfrost plus gold microscope slides were autoclaved with flexible silicon 12-well transwell adapters (Flexiperm, Starstedt), dried completely, and the transwell adapters mounted. Wells were coated with 0.1 mg/mL poly-l-lysine (0.1 M borate buffer pH 8.5) overnight at 37°C, aspirated, and dried. Cells were grown in the transwells overnight and subsequently treated with 10 μM UMB18 or 0.1% DMSO for 24h. At the end of the incubation, supernatants were aspirated, the transwell mount was removed, and the slides were submerged in ice-cold 50 mM ammonium formate (pH 6.7) for 1 minute with gentle rocking. Excess buffer was removed by tilting and using absorbent paper. Slides were dried, flat, in a desiccator for 20 minutes. Silver-assisted laser desorption ionization (Ag-LDI) was used to ionize cholesterol on a quadropole time of flight (qTOF) mass spectrometer in imaging mode (Yang2020_JMassSpectrom). Silver nitrate (8.5 mg/mL in 0.1% trifluoroacetic acid/methanol) was sprayed on the slides using an HTX M5 Matrix Sprayer with the following settings: 45°C nozzle, 40 mm height, 24 passes, 0.075 mL/min flow, 500 mm/min speed, 4 mm track spacing, 20 psi, 2 L/min N2, in “CC” pattern, with 0 s dry time between passes. Cholesterol imaging data were captured from an area of uniform monolayer (>80% confluence) integrity guided by an optical pre-scan using a Bruker Daltonics timsTOF Flex (Billerica, MA) in positive ion mode with a 50 μM scan area (11 μM x 11 μM beamscan), over range *m/z* 100-1000 with enhanced quadratic fit calibration to Agilent Tune-Mix, 500 shots per pixel at 10 kHz with 5-point setting in M5 small laser focus. Total scan area per sample was ∼3 mm^2^ resulting in ∼1200 pixels per replicate, with quadruplicate samples per condition. Data analyzed in SCiLS Lab (Bruker Daltonics), normalized (total ion current), projected with hotspotting on, and visualized as the ion *m/z* 493.260 +/-2.2 mDa ([cholesterol+Ag_107_]^+^) ion; the [cholesterol+Ag_109_]^+^ ion data agreed. Student’s T-test and a receiver-operator characteristic were performed between vehicle-treated and UMB18-treated cells on quadruplicate samples.

### Expression and purification of SKIC8

Transformed *E. coli* BL21 (DE3) cells containing the full-length human *SKIC8* gene with a HisX6 tag in the pET-28a(+) vector were grown in Luria-Bertani (LB)-Kanamycin broth at 37°C until the optical density at 600 nm equaled 0.8. SKIC8 protein expression was induced with 1 mM Isopropyl β-D-1-thiogalactopyranoside for 20 h at 15°C. Cells were lysed using Bugbuster (Novagen) supplemented with protease and phosphatase inhibitors (Thermo Scientific). Lysates were clarified by centrifugation at 14,000 rpm for 30 min. and the supernatant was applied to a Talon Metal Affinity Resin (Clontech) column equilibrated in 50 mM sodium phosphate (pH 7.8) containing 10 mM imidazole and 300 mM NaCl. The column was washed with the same sodium phosphate buffer containing 20 mM imidazole. SKIC8 was eluted in 50 mM sodium phosphate (pH 7.8) containing a 50-250 mM imidazole gradient and 300 mM NaCl. The fractions containing SKIC8, as determined following SDS-PAGE, were pooled, and concentrated with an Amicon ultrafiltration cell (Millipore, Bedford, MA). The concentrated protein was dialyzed against 20 mM Tris-HCl (pH 7.8) containing 150 mM NaCl.

### Surface Plasmon Resonance (SPR)

SKIC8 and test compound interactions were evaluated using a Biacore T-200 instrument (GE Healthcare). A carboxymethylated dextran chip (CM5, GE Healthcare) was used to immobilize SKIC8. A net positive charge is required for direct amine coupling; therefore, the proteins were diluted to a final concentration of 30 µg/ml in 10 mM sodium acetate buffer pH 4.0. The protein was injected at 2 µl/min until approximately a 4500 resonance/response unit level of immobilized SKIC8 was reached, and then the flow cell was deactivated with 1 M ethanolamine HCl (pH 8.5). Degassed 20 mM Hepes (pH 7.4) containing 5 mM MgCl2, 100 mM NaCl, and 0.005% Nonidet P40 was utilized as continuous running buffer for all experiments. Test compound binding interactions with SKIC8 were investigated by injecting test compound (1.0 –50 µM) at a flow rate of 20 µl/min. The resulting sensorgrams were analyzed using BIAevaluation 3.1 software (GE Healthcare) to determine the dissociation constant (K_D_) as previously described(32).

### RNAi knockdown

Cells were seeded to 24-well plates at a density of 4.4e4 cells/well one day prior to transfection. Transfections were performed using Oligofectamine (Thermo Scientific) to the manufacture’s methods. Briefly, per well of transfection, 4.4μl Opti-MEM (Gibco) was mixed with 2.2μl Oligofectamine and incubated at room temperature (RT°C) for 5 min, while 35.5μl Opti-MEM was mixed with 0.8μl 50mM siRNA in a separate tube. The two tubes were combined and incubated at RT°C for 20 minutes prior to addition of 177μl Opti-MEM. Cells were washed 1x with PBS and 1x with OptiMEM then 200μl of transfection mixture was added per well. All siRNAs were purchased from Sigma using their Rosetta prediction system for predesigned sequences. MISSION siRNA Universal Negative Control #1 (Sigma) was used as the scrambled control sequence. Sequences are displayed in table:

#### RNAi sequences

| Gene name | siRNA ID |
| --- | --- |
| **SKIC2 (A.K.A SKIV2L)** | SASI_Hs01_00196881 |
| **SKIC3 (A.K.A TTC37)** | SASI_Hs01_00010354 |
| **SKIC8 (A.K.A WDR61)** | SASI_Hs01_00155941 |
| **SCAP si-1** | SASI_Hs01_00032110 |
| **SCAP si-2** | SASI_Hs01_00032111 |
| **SREBF1 si-1** | SASI_Hs01_00051828 |
| **SREBF1 si-2** | SASI_Hs01_00051829 |
| **SREBF2 si-1** | SASI_Hs01_00075424 |
| **SREBF2 si-2** | SASI_Hs01_00075425 |
| **SREBF2 si-3** | SASI_Hs01_00075426 |

### Western blotting

Triton-X100 (Tx100) lysis buffer lysis buffer (1% Tx100 (v/v), 150mM NaCl, 50mM Tris (all Sigma), pH8). In all cases 1x cOmplete Mini, EDTA-Free protease inhibitor cocktail (Roche) was added to the lysis buffer and for experiments involving S6K and phospho-S6K (pS6K), Phosphatase Inhibitor Cocktail 2 (Sigma) was additionally used. Western blotting procedures have been previously described(33). Primary antibodies used as follows: p70 S6K (Cell Signaling Technologies #9202S, 1:1000 dilution), phospho-p70 S6K (T421/S424) (Cell Signaling Technologies #9204S, 1:1000 dilution) and mouse anti-tubulin (clone DMA1A, Sigma, 1:1000). Secondary antibodies were used as follows: goat anti-rabbit HRP and goat anti-mouse HRP (both 0.8 mg/ml, Thermo Scientific, 1:10,000 diluted) which were detected using ECL Prime (Amersham).

### Data analysis

All graphs produced and statistical analysis performed using GraphPad Prism 10.5 software.

## Acknowledgements

We would like to thank all of the Frieman, Jackson and Coughlan lab members for their help in this study.

## Author Contributions

- Conceptualization: SW, MBF
- Data curation: LT,
- Formal analysis: SW, LT, AS, PS,
- Funding acquisition: MBF
- Investigation: SW, LB, AS, GK,
- Methodology: SW, LT, AS, PS,
- Resources: ADM
- Supervision: MBF
- Visualization: SW, LT, AS, PS,
- Writing – original draft: SW, AS, PS, MBF
- Writing – review & editing: SW, MFB

## Patents

Filed patents: Patents on UMB18 and related compounds have been filed through University of Maryland, Baltimore and listed as PCT/US20/36483 and PCT/US2021/1061863.

## Data and Material Availability

All data needed to evaluate the conclusions in this paper are present in the paper and supplementary materials. Data are also deposited at https://osf.io/63h8t/.

